# Tumor cell intrinsic RIG-I activation co-opts the host microenvironment to drive immune mediated tumor rejection

**DOI:** 10.1101/2022.12.21.521495

**Authors:** Eugenia Fraile-Bethencourt, Sokchea Khou, Adrian Baris, Rebecca Ruhl, Sudarshan Anand

## Abstract

Targeting cytosolic nucleic acid sensors is a potent approach to drive type I interferon responses and anti-tumor immunity. Recent evidence suggests that activation of retinoic acid inducible gene-I (RIG-I) using synthetic hairpin RNA agonists decreases tumor progression in multiple preclinical models. However, the role of tumor cell intrinsic RIG-I in shaping tumor cell fates and the host immune microenvironment remains unclear. Here, we show that RIG-I expression is correlated with better overall survival and a distinct immune gene signature in specific human cancers including colorectal cancer. Activation of RIG-I in breast and colorectal cancer cells is sufficient to drive tumor cell death in vitro and significantly delay tumor growth in vivo in multiple preclinical models. Importantly, the efficacy of tumor cell RIG-I activation is lost in immune deficient mice suggesting the requirement of immune responses for this effect. We observe that tumor cell intrinsic RIG-I activation elicits a robust cellular and molecular immune response. We show that tumor cell RIG-I activation also leads to induction of specific immune checkpoints including PD-L1. Using a publicly available database, we found that RIG-I expression serves as an excellent prognostic marker for responders to checkpoint immunotherapy, particularly PD-L1/PD-1 across cancers. Finally, combination of tumor cell intrinsic RIG-I activation with anti-PD-L1 led to a synergistic decrease in tumor growth in a colorectal tumor model. Our findings suggest that tumor cell intrinsic RIG-I can be targeted to enhance anti-tumor immune responses and highlights a potential strategy for anti-cancer vaccines that can invigorate the immune system.

## Introduction

The tumor microenvironment (TME) plays a critical role in cancer progression [1, 2]. One emergent strategy in promoting an anti-tumor microenvironment is the activation of the innate immune response by activating the cytosolic nucleic acid sensors in the TME [3]. The nucleic acid sensors are pattern recognition receptors (PRRs) that recognize DNA or RNA from pathogens. Among the different nucleic acid sensors, the RNA sensor the retinoic acid-inducible gene I (RIG-I) is widely express across cell types, including cancer cells. RIG-I is part of the RIG like receptors (RLR) family, and it is encoded by the DDX58 gene. RIG-I recognizes 5’di or tri-phosphorylated short double strand RNA (dsRNA) in the cytosol. Upon activation, RIG-I dimerizes and complex with the mitochondria antiviral signaling protein (MAVS), triggering a rapid and potent anti-viral response via NF-kB and IRF3/7. Thus, activation of RIG-I leads to IFN-I gene expression as well as to release of proinflammatory cytokines. Moreover, this response induces an apoptosis called RLR-induced IRF3 mediated pathway of apoptosis (RIPA), in which IRF3 is not only translocated into the nucleus, but also recruited by Bax to the mitochondria, where IRF3 triggers the apoptosis via cytochrome c release [4-6].

Emerging new studies show that activation of RIG-I augments the immune response in cancer. For instance, RIG-I activation in pancreatic cancer cells drives a program of immunogenic cell death [7]. RIG agonists increase responsiveness to checkpoint blockade in preclinical models of melanoma and leukemia [8-10]. These studies identify significant cellular and molecular immune correlates of RIG-I inflamed tumors that correspond to a good anti-tumor response. Interestingly, intra-tumoral injections of RIG-I agonists also decrease tumor burden in preclinical models and paved the way for a phase I trial in solid tumors. The phase 1 trial with 20 patients showed that the RIG-I agonist drug was well tolerated and demonstrated preliminary evidence of clinical efficacy with shrinkage of injected and non-injected lesions [11]. These observations suggest that simultaneous activation of the innate and adaptive arms of the immune system by activating RIG-I may generate therapeutic responses across several cancer types and this is a feasible strategy in human cancer [12, 13].

While these studies highlight the potential of targeting this pathway in cancers, it is not clear if the activation of RIG-I in the tumor cells or the TME or both are responsible for the biological effects on tumor growth and immune activation within tumors. Here, we show that tumor cell intrinsic RIG-I activation is sufficient to increase tumor cell death, diminish tumor growth and activate cellular and molecular correlates of anti-tumor immune responses in preclinical models of breast and colorectal cancers. Our results demonstrate common RIG-I activation signatures across cancer types in human cancers and identify gene signatures and pathways that are activated by RIG-I in breast and colorectal cancer. Finally, our data showcases that RIG-I expression correlates with patient responses to immune checkpoint blockade, particularly PD-L1/PD-1 antagonists and demonstrates the functional significance of combining tumor cell RIG-I activation with immune checkpoint blockade. These observations highlight an underappreciated role for RIG-I within the tumor cell compartment and highlight potential opportunities for rational combinations of immune targeted agents with RIG-I activation.

## Materials and Methods

### Cell culture

MC-38 (Kerafast #ENH204-FP) cell line was cultured in Dulbecco’s Modified Eagles Medium (DMEM), High Glucose (HyClone™) supplemented with 10% Fetal Bovine Serum (FBS) (Bio-Techne R&D Systems #S11550H). CT-26 (ATCC #CRL-2638), 4T1 (ATCC #CRL-2539) and E0771 (CH3 Biosystems #940001) cell lines were culture in RPMI 1640 with L-Glutamine (Lonza™ BioWhittaker™ # BW12115F12) supplemented with 10% of FBS. Each cell line was tested and found negative for mycoplasma contamination before use in the assays described.

### Transfection

Approximately 1×10^5^ cells were seeded in 6 well plates, incubated 24h and transfected at 50– 60% confluence. RIG-I agonist (Invivogen #tlrl-hprna) or control agonist (Invivogen #tlrl-3prnac) were transfected with a cationic lipid-based transfection reagent -LyoVec^TM^ (Invivogen #lyec-12) following tmanufacturer’s instructions.

### Cell Titer-Glo/Caspase Glo

Cells were transfected with RIG-I or control agonist in white 96 well plates (Greiner Bio-One™ CellStar™). Viability, survival was assessed using Cell Titer-Glo and Caspase 3/7 Glo assays (Promega) and analyzed at 24–48h post-treatments, according to manufacturer’s instructions.

### Clonogenic survival assay

Cells were seeded in 6 well plates at low density (approximately 500 cells per well) and treated with RIG-I or control agonist. Plates were incubated for 5-7 days before staining. Cells were wash with PBS and fixed using a solution of paraformaldehyde 4% and crystal violet 0.5% during 15 min in a shaker. Cells were washed and colonies were counted using ImageJ software.

### Western blot and Simple Wes

Cell lysates were prepared by harvesting cells in RIPA buffer (Pierce #89900) containing Protease Inhibitor Mini Tablets (1/10 mL RIPA buffer, Cat: A322953, Pierce) with phosphatase inhibitor cocktail 2 and 3 (1:1000, Cat: P5726 and P0044) and centrifugation at 4°C. Lysates from the tumors were obtained by homogenization in a handheld tissue homogenizer following the addition of RIPA buffer and centrifugation as described before. Lysates were quantified using a BCA assay (Pierce, #23227) kit. For western blot, equal amounts of protein were loaded on a 4–12% gradient SDS-polyacrylamide gel (NuPAGE, Life-Technologies) and transferred onto polyvinylidene difluoride (PVDF) membranes. Membranes were blocked Intercept® (TBS) Blocking buffer (Li-cor #P/N 927-60001) and incubated with the antibodies named in the following table. Membranes were washed in TBST and incubated with secondary antibodies from Licor Biosciences were used goat anti mouse 925-68020 (1:15,000) and goat anti rabbit 925-32211 (1:15,000). Blots were scanned on the Licor Odyssey scanner according to manufacturer’s instructions. For Simple Wes, samples were diluted with 1× Sample Buffer (ProteinSimple). Protein quantification was performed using a 12–230 kDa 25 lane plate (Cat: PS-MK15; ProteinSimple) in a ProteinSimple Wes Capillary Western Blot analyzer according to the manufacturer!s instructions.

For specific protein array experiments with membrane arrays-mouse XL cytokine Array (R&D Biosystems-ARY028) protocol were according to manufacturer’s instructions.

| Antibody | Company | ID | Host specie | kDa |
| --- | --- | --- | --- | --- |
| RIG-I | CellSignal | 3743S | Rabbit | 102 |
| NF-kB | Abcam | ab32536 | Rabbit | 65 |
| STAT1 | CellSignal | 14994T | Rabbit | 84 |
| ISG15 | CellSignal | 2743S | Rabbit | 16 |
| GAPDH | CellSignal | 5174S | Rabbit | 37 |

### RNA extraction, RT-PCR and gene expression

Genomic RNA from cells and homogenized tumors was extracted using the GeneMATRIX Universal RNA Purification Kit (EuRx #E3598) following the manufacturer!s instruction. Samples were treated with DNAse I (Invitrogen™ #18068015) to avoid DNA contamination. Reverse transcription was performed with 500ng of RNA using the High-Capacity cDNA Reverse Transcription Kit (Cat: 4368814, Applied Biosystems). Gene expression was assayed by quantitative PCR using TaqMan^TM^ Master Mix II no UNG (Cat 4440049, Thermofisher Scientific) with the following probes: Ddx58 (Mm01216853_m1), Mx1 (Mm00487796_m1), Isg15 (Mm01705338_s1), Ifi44 (Mm00505670_m1), Oas1 (Mm04209919_m1), Gapdh (Mm99999915_g1). Data was normalized to internal control Gapdh and gene expression quantified using the using the 2^−ΔΔCt^ method with the control treatment as reference.

### RNA sequencing

RNA from treated cells was extracted as previous mentioned. Samples were quantified and processed by Massively Parallel Sequencing Shared Resource at OHSU. Sequencing was done on either a HiSeq 2500 or a NovaSeq 6000. Base calling was done using RTA 1.18.66.3 (HiSeq) or RTA 2.11.3 (NovaSeq). RNA-Seq library preparation with stranded with poly(A)+ selection by Illumina following the manufacturer!s instructions. Normalization and DESeq2 analysis was done using a web-based software RaNAseq (https://ranaseq.eu/index.php).

### ELISA

Mouse IFN-β was measured using Quantikine®ELISA (R&D Systems #MIFNB0) following the manufacturer!s instructions. Supernatant from control and RIG-I activated cells was collected 24h after treatment. Control samples were directly assayed while RIG-I activated sampled were diluted 1:2 in RD1W buffer.

### In vivo assays

Animal work was approved by the OHSU Institutional Animal Use and Care Committee. Protocols were performed in accordance with the relevant guidelines and regulations. All 8–10- week-old C57BL6 or Balb/C mice were purchased from Jackson Labs. Mice were injected in two mammary fat-pads with 1×10^5^ 4T1 or E0771 cells or subcutaneously in each flank with 1×10^5^ CT26 or MC38 cells. Cells were prepared in 50% of Growth Factor Reduced Matrigel (Fisher-Scientific #CB-40230C) for injection of 50-100 µl per mouse. Tumor growth was measured with digital calipers with volume computed as ½ × Length × Width^2^. For treatments, mice were randomized into groups when tumor volume reached 80–120 mm^3^, approximately 6-8 days after tumor cell injection. 100 μg of anti-PD-L1 (10F.9G2, BioXcell) or isotype control antibody (2A3, BioXcell) was injected i.p on indicated days. Treatment personnel were blinded to the identity of the groups wherever feasible. Tumor weights were measured at endpoints.

### Tumor digestion

Once tumors reached the appropriate size/endpoint, tumors were harvested and digested. Mechanically disaggregated tumors were enzymatically digested using a solution of 1mg/ml of collagenase IV (Sigma-Aldrich #C5138) and 200µg/ml of DNase I (Sigma-Aldrich #11284932001) in DMEM for 20min at 37°C in a shaking platform. To stop the reaction, tubes were placed on ice. Tumor cells were filtered using 40µm nylon strainers and FACs buffer. Finally, cells viability and cell number were calculated using Beckman Optima TLX cell counter.

### Multicolor Flowcytometry

Cells were plated on a 96-well plate (2 × 106 cells per well), blocked with Fc-Block (BD Bioscience, cat # 553142). 1:200 and Live/ Death Aqua reagent solution 1:500 for 25 min on ice. Subsequently cells were stained with an antibody cocktail mix containing the following antibodies with the indicated fluorophores. After staining, cells were fixed with Cytofix Buffer (BD Bioscience, cat # 554655). After the staining cells, were washed and resuspended in 200 ul of FACS buffer and stored at 4 °C protected from light until analysis in a BD-LSR-Fortessa cell analyzer. Data was analyzed with FlowJo v10.8.0 software.

**Table 1:**
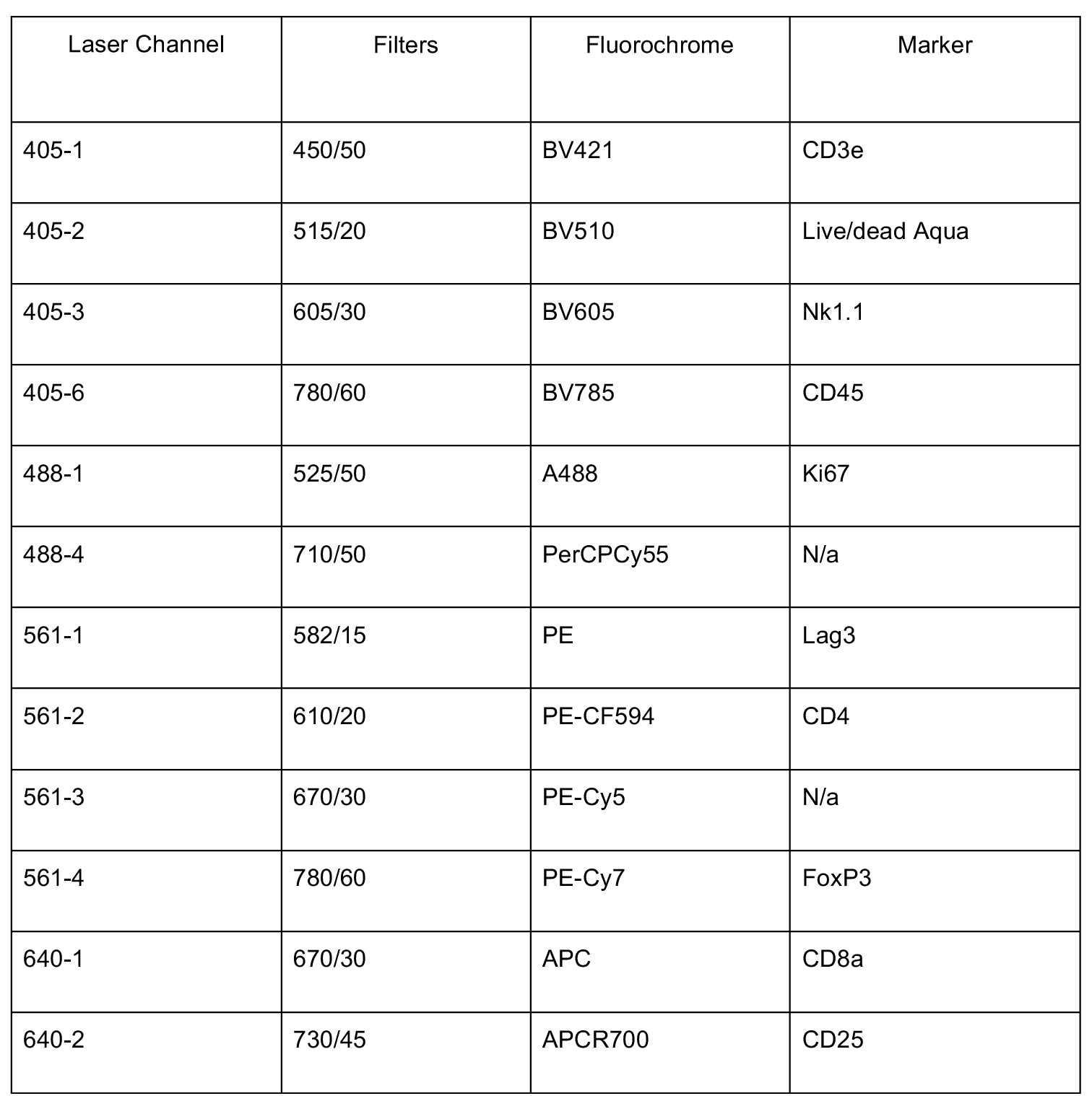
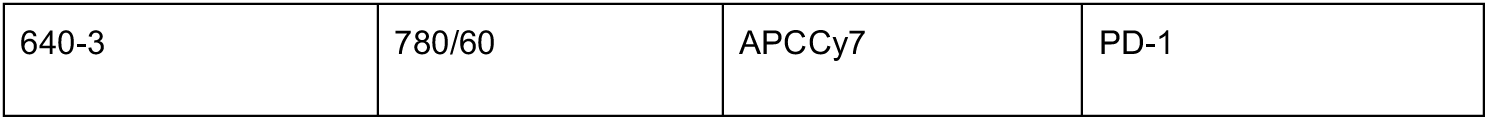
Myeloid flow immune panel markers.

**Table 2:**
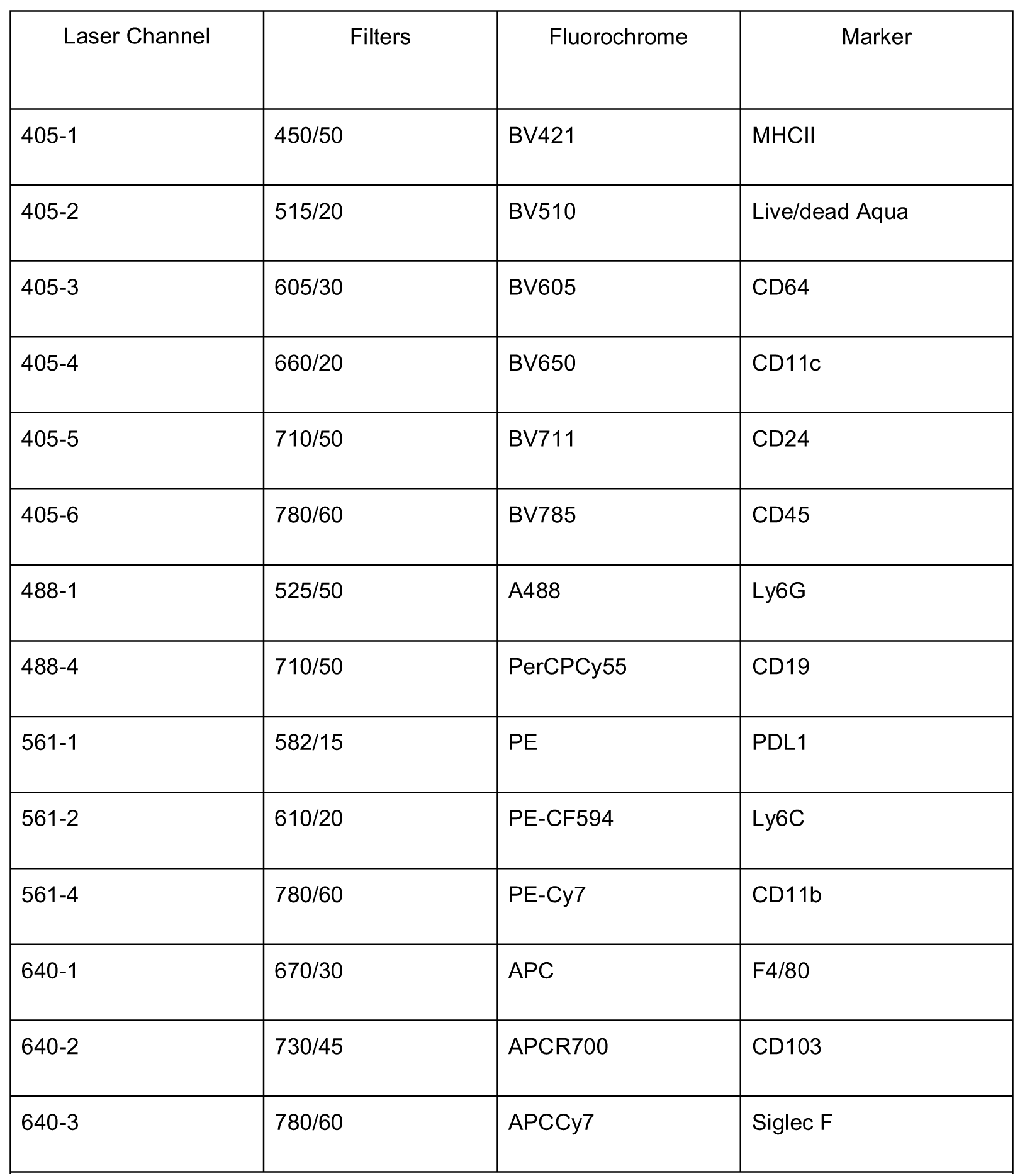
Lymphoid flow panel immune marerks.

### NanoString

NanoString immune profiling was performed using nCounter® PanCancer Immune Profiling Panel (NanoString XT_PGX_MmV1_CancerImm_CSO #115000142. Samples were hybridized overnight at 65°C according to the manufacturer!s instructions. The cartridges were run in the NanoString nCounter SPRINT Profiler and data was analyzed using nSolver Analysis Software 4.0. Immune signatures were analyzed using the advanced analysis module of the nSolver Software.

### Statistics

All statistical analysis was performed using Prism software (GraphPad Software, San Diego, CA). Two-tailed Student!s T-test or one-way ANOVA with post-hoc (Tukey!s or Holm-Sidak!s) corrections was used to calculate statistical significance. Two-tailed Student!s T-test was used to calculate statistical significance when groups were normally distributed. For more than two groups, two-way ANOVA was used. For data that were not normally distributed, Mann– Whitney U-test was used. Variance was similar between treatment groups. For experiments where the data was not normally distributed, we used the Kruskal-Wallis test. P-values <0.05 were considered significant.

### Data and Material Availability Statement

All RNA sequencing data has been deposited to Gene Expression Omnibus GEO. Request for materials should be addressed to the corresponding author.

## Results

### RIG is expressed in human cancer and is associated with a complex immune microenvironment

We probed the expression of RIG across human cancers in the TCGA. In a majority of cancers, we found RIG-I mRNA levels were upregulated in a subset of patients (Fig 1A). For some cancers – breast cancer, kidney renal cell carcinomas, lung adenocarcinomas, ovarian cancer, rectal adenocarcinomas and sarcomas, expression of RIG-I was associated with an increase in overall survival (Fig 1B). Using an immune microenvironment analysis tool TIMER (25), we set out to investigate whether there was a correlation between RIG-I and immune responses. We found RIG-I expression correlates with expression of CD8, perforin and granzyme expression (Fig 1C) gene signatures that are typical of robust anti-tumor immune responses. Interestingly, RIG-I expression also correlates with the expression of several immune checkpoints including CD80, CD86, PD-L1, CTLA4, and LAG3 (Fig 1C). We compared the immune microenvironment of RIG-I high vs RIG-low CRCs from a cohort of 121 patients and observed a distinct type I interferon signature (Fig 1D). Importantly, the molecular immune signatures in our cohort were validated in the TCGA COAD dataset (Fig 1F). These observations indicate that expression of RIG-I itself correlated with better survival in a few cancers and the molecular signatures suggest a type-I interferon response gene signature indicative of RIG-I activity.

**Figure 1.**
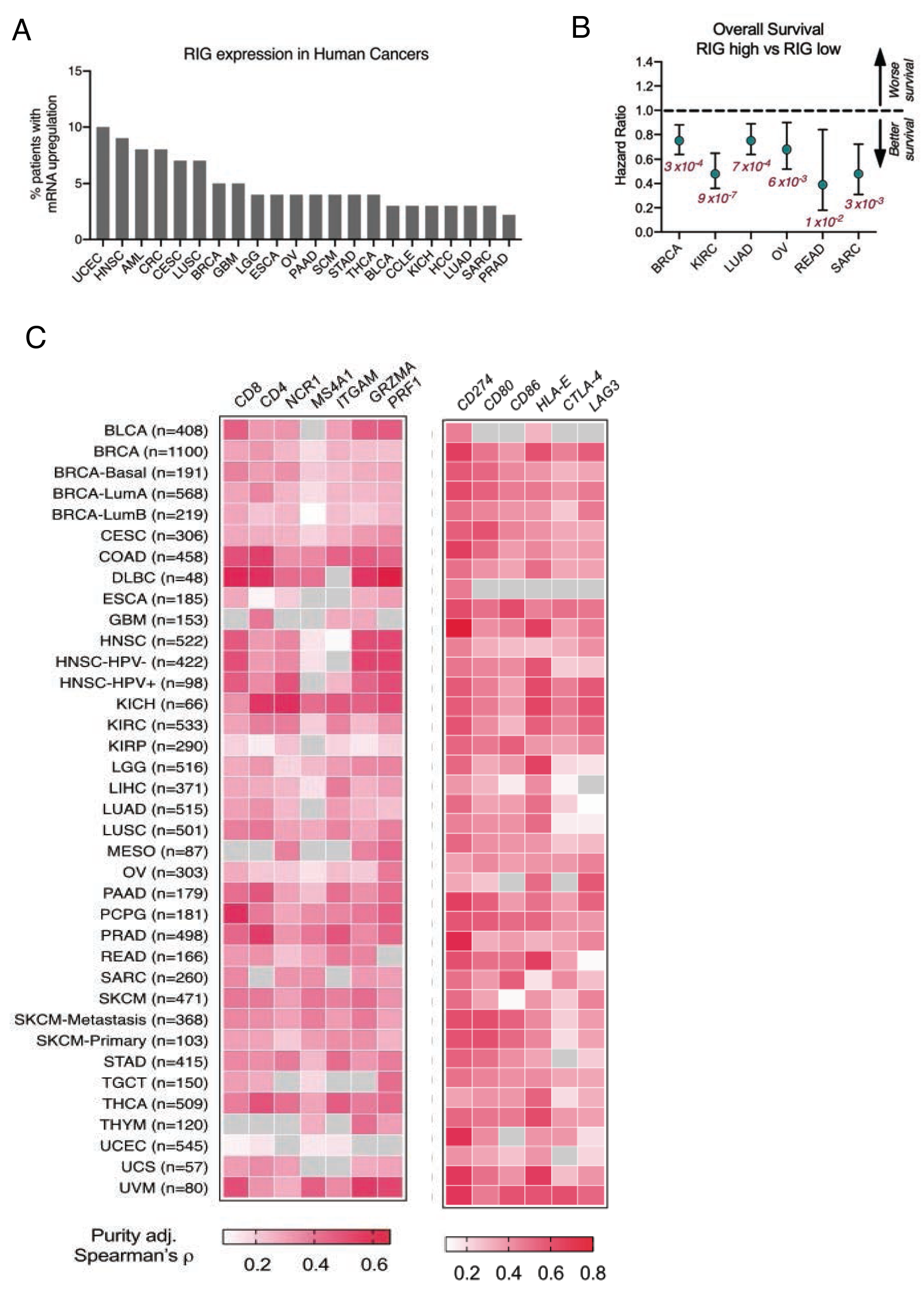
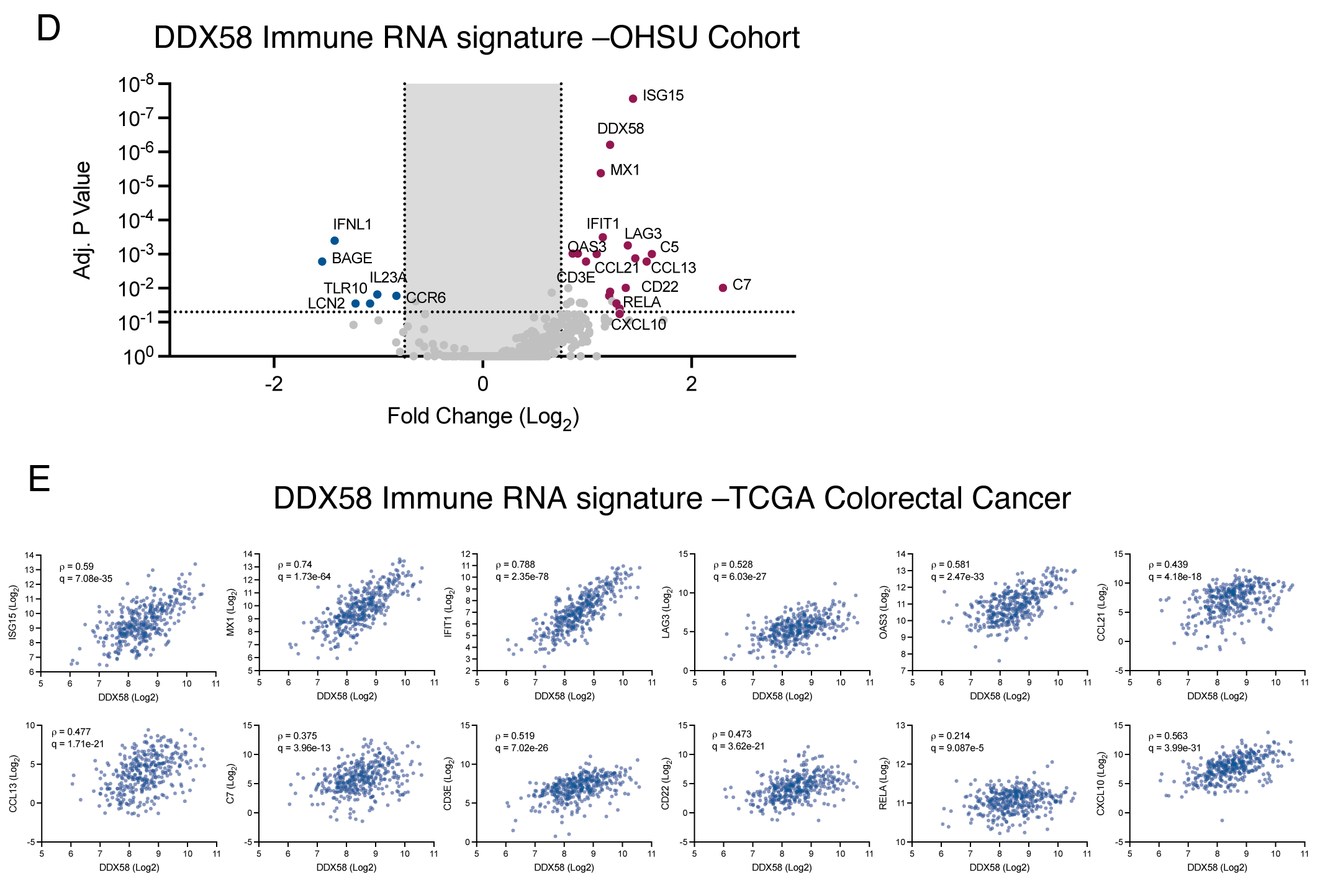
RIG-I expression correlates with survival and immune contexture in human cancer. Percentage of patients with mRNA upregulation of RIG1 gene DDX58 across cancer types. Data from TCGA datasets accessed via cBioportal. B) Overall survival of patients with high or low RIG expression across cancer types. Survival data was analyzed using Kmplot datasets with patients stratified by median expression level into high vs low groups. Error bars show 95% confidence intervals. Log-rank P values in magenta. C) Correlation coefficients of immune cell markers with RIG-I gene expression across TCGA. Data was analyzed using TIMER portal. Grey squares = correlations that were not statistically significant. D-E) DDX58 associated immune signature in CRC - OHSU cohort.

### RIG-I activation drives a potent type I interferon response and increases tumor cell death

To evaluate a causal relationship between RIG-I expression and tumor burden, we performed a series of *in vitro* and *in vivo* experiments. We used a commercially available RIG-I agonist, 5 ! triphosphate stem loop RNA derived from the H1N1 influenza virus (Supplementary Figure 1A) and a commercial dsRNA without the triphosphate as a control agonist. We confirmed that RIG-I agonist drove an interferon response in CT26 murine colorectal carcinoma cells and E0771 mammary carcinoma cells by evaluating several ISGs including RIG itself and phospho-NF-κB, STAT1 and ISG15 (Supplementary Figure 1B). We confirmed that this agonist was also able to induce type-I interferon robustly across different breast and colorectal cancer cell lines (Supplementary Figure 1C). Transfection of our RIG-I agonist decreased proliferation (data not shown), increased apoptosis (Figure 2A) and decreased colony formation in a clonogenic survival assay in multiple colorectal, breast cancer cell lines (Figure 2B-C). We then performed RNA sequencing on CT26, MC38 and E0771 cell lines at early (24h) and late (72h) time points after treatment (Figure 3A and Supplementary Figure 2) and identified a set of common RIG-I activation induced genes across these cells. Pathway analysis using Enrichr [14, 15] identified a group of most responsive pathways that were slightly different across different cell lines (Figure 3B-D) but had common pathways such as interferon response and apoptosis that are consistent with our phenotypic readouts. Finally, we compared RIG-I correlated genes in the human COAD dataset with our RNAseq data from murine CRC cell lines treated with the RIG-I agonist and found a 37 gene RIG-I activation signature specific for CRC (Figure 3E).

**Figure 2.**
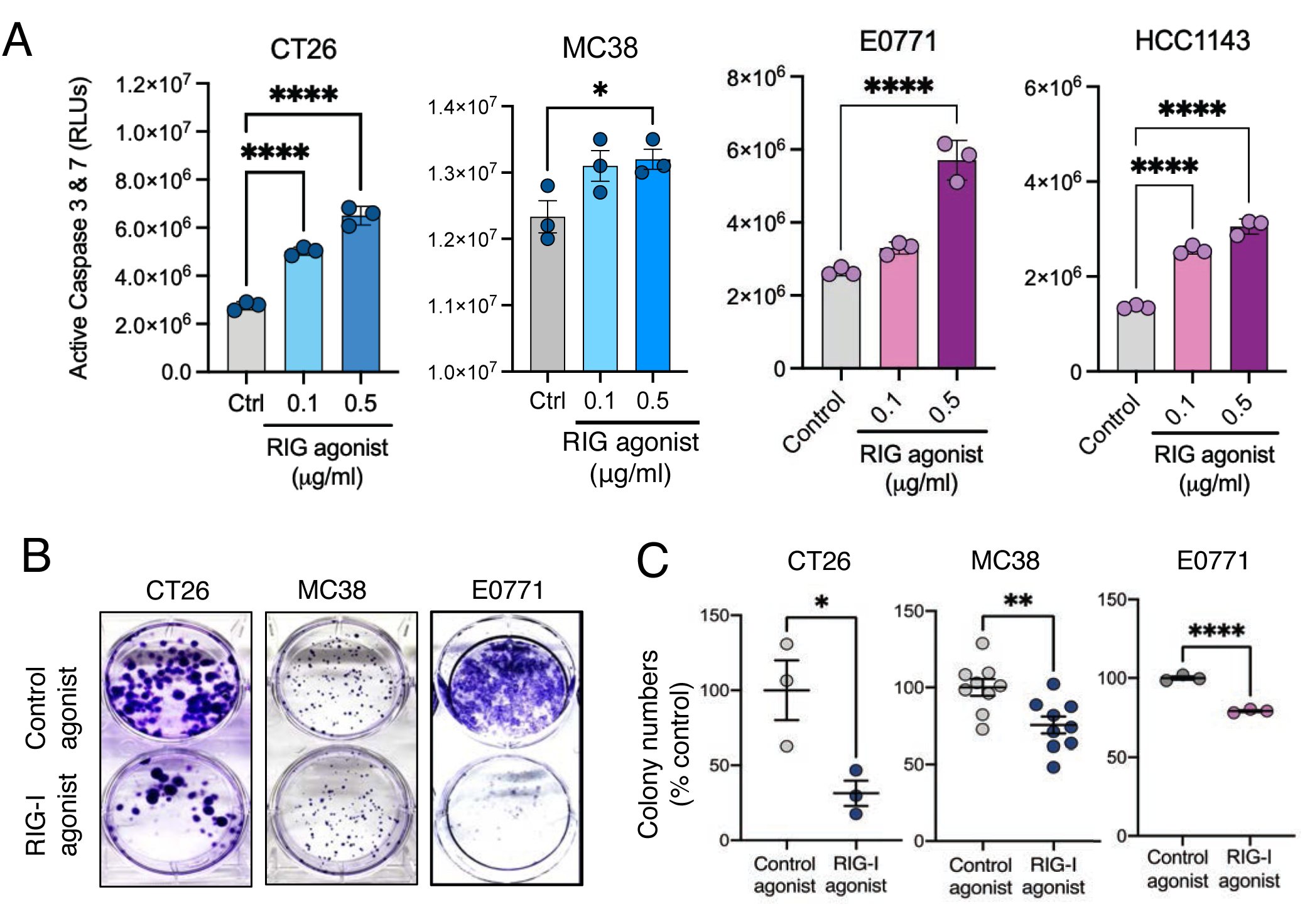
Tumor cell RIG-I activation causes apoptosis. A) Active Caspase 3 & 7 as measured by a CasGlo assay in the indicated cell lines treated with Control agonist or RIG-I agonist. B-C) Clonogenic survival assay in the indicated cell lines 7-12d after control or RIG agonist treatment. P<0.05, ** P<0.01, *** P<0.001, ****P<0.0001 using ANOVA with post-hoc Sidak’s test or a Student’s T-test for comparison across two sample groups.

**Figure 3.**
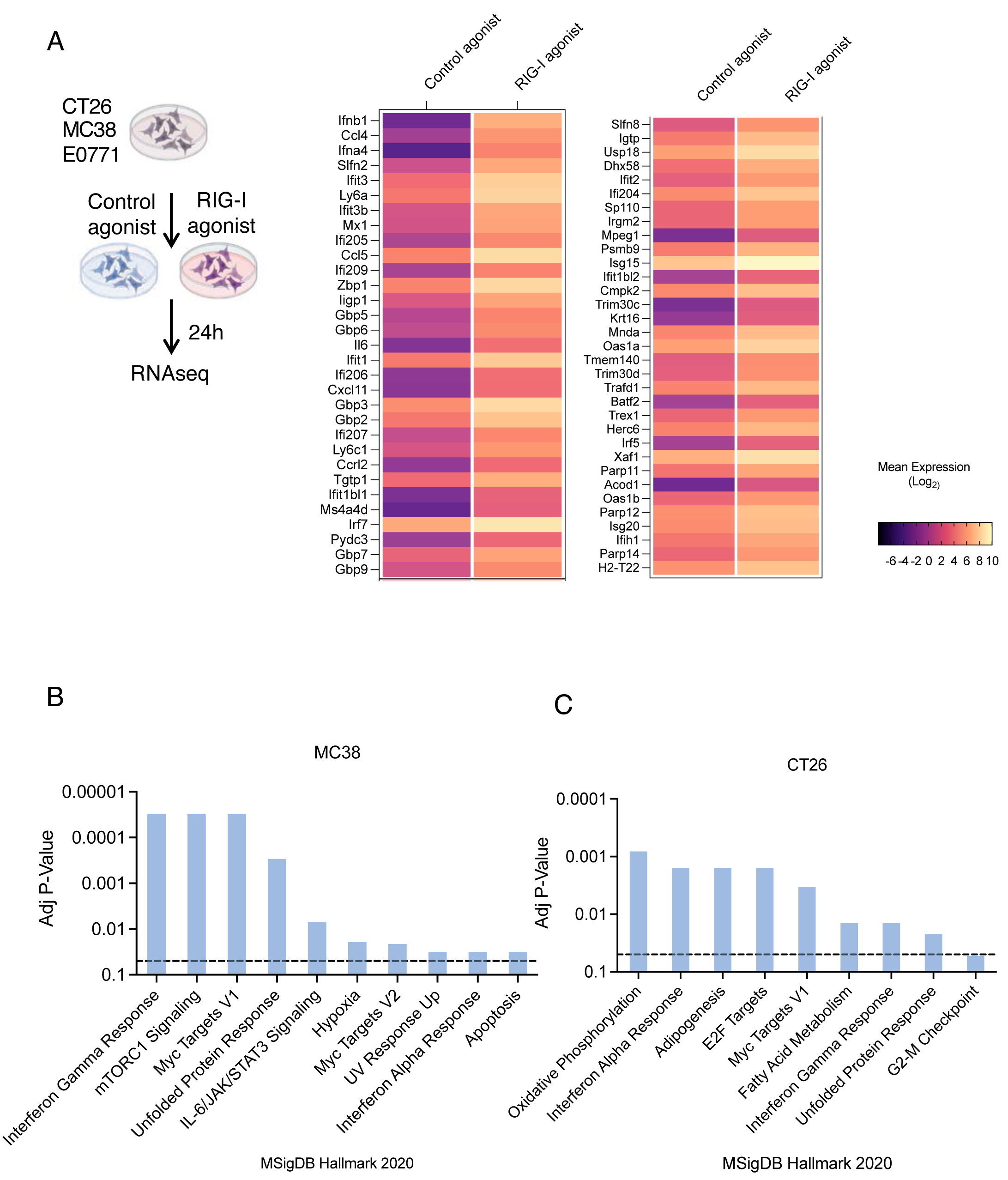
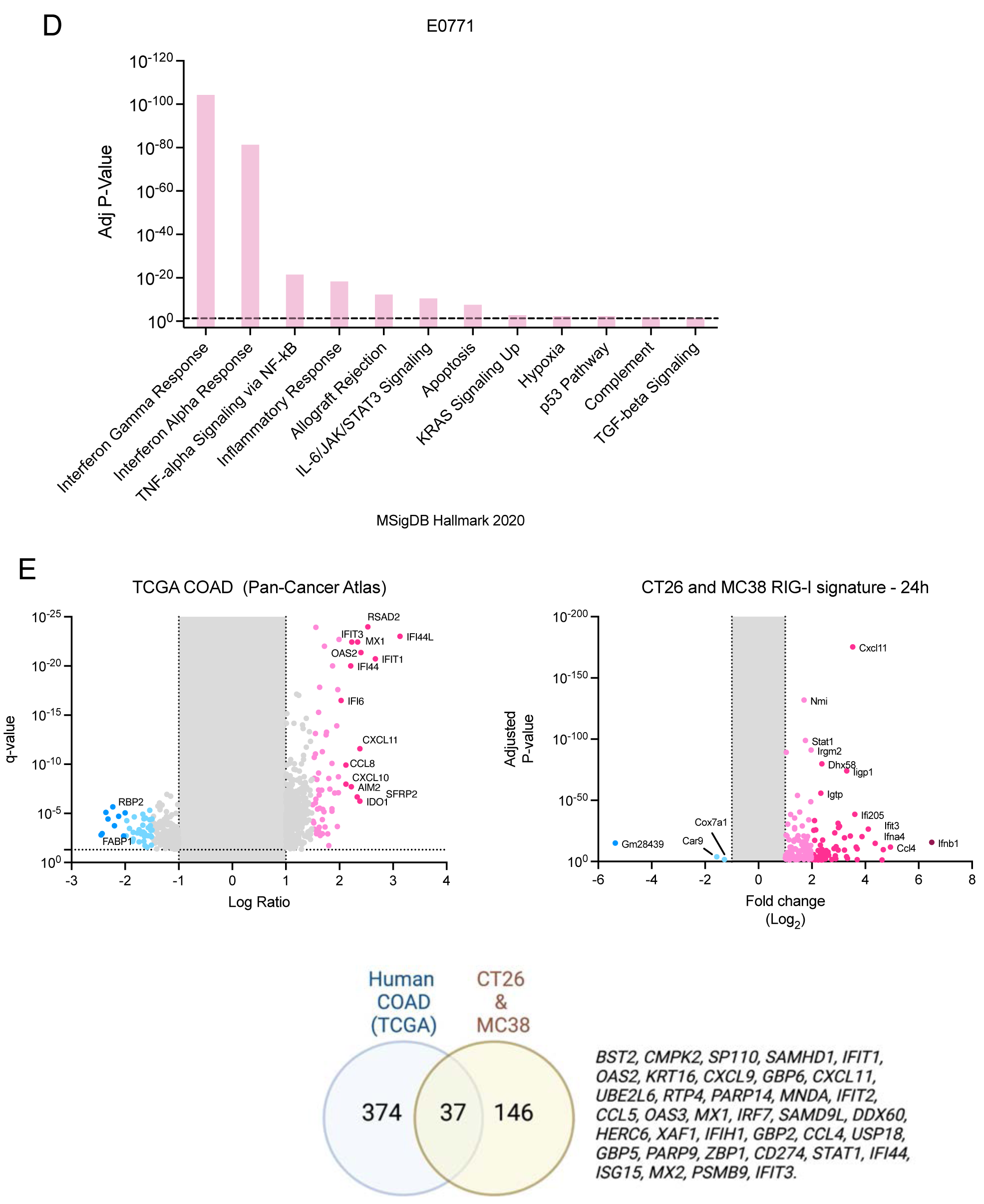
Tumor intrinsic RIG-I activation enhances immune gene expression. A) Heatmap depicting significant (>4 fold, adj. p-value <0.05) RIG-I agonist induced gene expression changes across three tumor cell lines. B-D) Gene Set Enrichment Analysis showing significantly impacted Hallmark pathways in each of the tumor cell lines. E) Volcano plots depicting RIG-I correlations in TCGA –COAD dataset (Pan-Cancer Atlas) and RNAseq data from mouse CRC cell lines 24h after RIG-I agonist treatment. Venn diagram depicts the 37 gene RIG-I signature in common between the human and mouse datasets.

### Tumor cell intrinsic RIG-I activation slows tumor growth

We next tested if RIG-I activation in tumor cells was sufficient to decrease tumor growth in preclinical models. In two different mouse tumor models-subcutaneous MC38 and CT26 colorectal carcinomas and orthotopic E0771 mammary carcinomas in C57BL/6 mice, we observed tumor cell RIG-I activation alone is sufficient to decrease tumor burden (Fig 4 A-C). In contrast to the immune competent models, RIG agonist transfected tumors grew at the same rate as control agonist transfected tumors in NOD-SCID-γ chain knockout (NSG) mice indicating that the immune response is critical in decreasing tumor burden downstream of RIG-I activation (Fig 4D).

**Figure 4.**
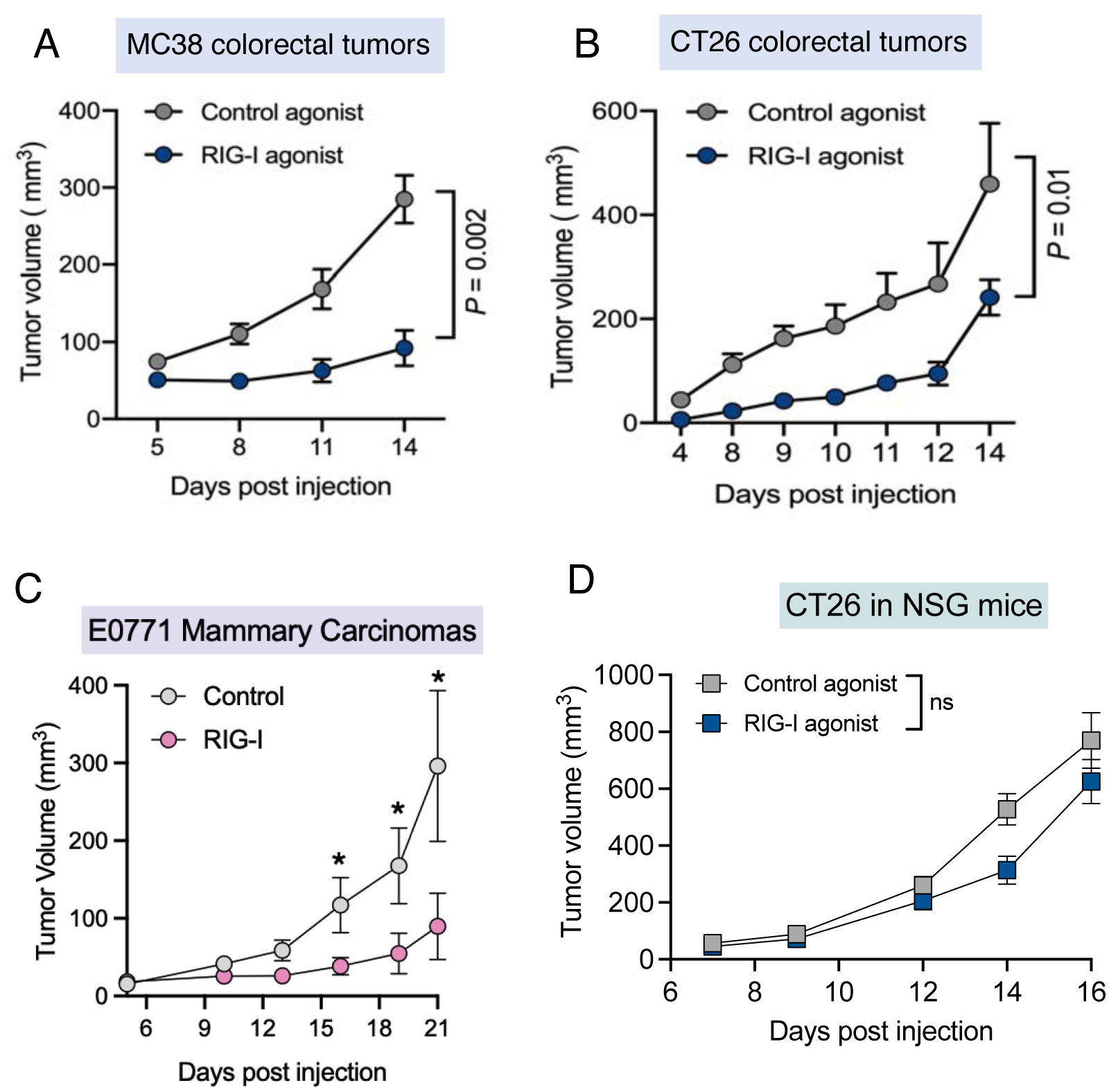
Tumor cell intrinsic RIG-I activation decreases tumor growth in immune competent but not immunocompromised mice. Control agonist or RIG-I agonist transfected MC38 (A), CT26 (B) murine colorectal carcinoma cells or E0771 (C) mammary carcinoma cells were injected in C57BL/6, Balb/C mice and C57BL/6 mice (n=5 per group) or D) NOD-SCID ! chain knockout (NSG) mice and tumor growth was monitored using calipers on the indicated days. Tumor growth P values from two-way ANOVA with post-hoc Sidak’s correction or Mann-Whitney U-test (C).

### Tumor cell RIG-I expression enhances molecular and cellular immune responses

We compared the immune microenvironment of RIG-I inflamed tumors in the MC38, CT26 colorectal carcinomas using Nanostring RNA profiling and multi-color flow cytometry (Supplementary Figure 3). We found that RIG-I activated tumors induced a strong type I interferon program *in vivo* (Figure 5A-B). At the cellular level, we observed that RIG-I activation in tumor cells attracted more NK cells, dendritic cells and CD4 T-cells across both MC38 and CT26 models (Figure 5C-D) and decreased CD8 T-cell exhaustion as measured by PD-1, LAG3 expression in the CT26 model (Figure 5D). Interestingly, we found that tumor intrinsic RIG-I activation is sufficient to diminish tumor growth measurably in athymic nude mice (Supplementary Figure 4) with a significant increase in NK cell infiltration. These data indicate that the activation of RIG-I in tumor cells is sufficient to enhance the cellular and molecular immune responses in the tumors and NK cell activity maybe critical for the anti-tumor responses.

**Figure 5.**
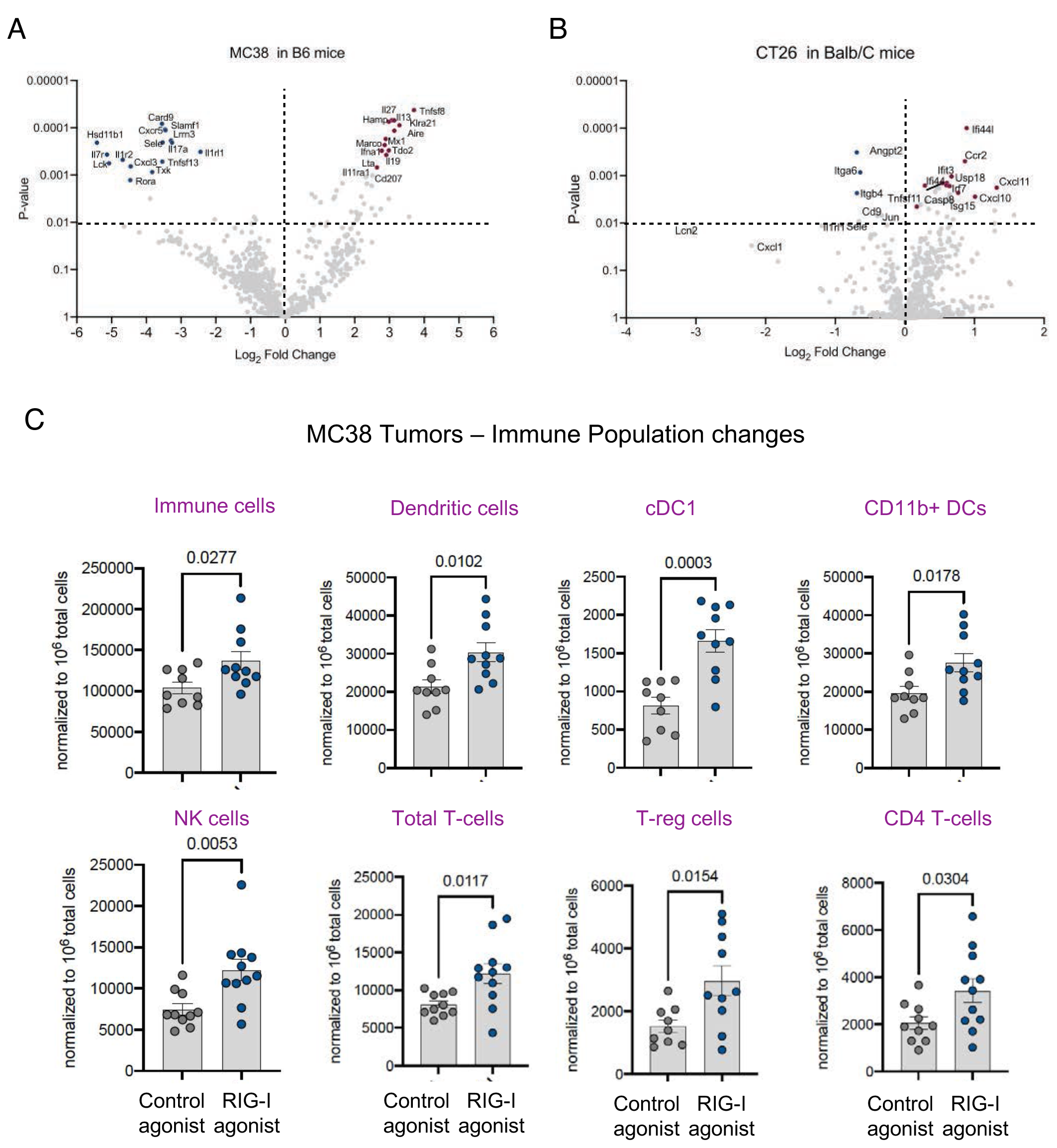
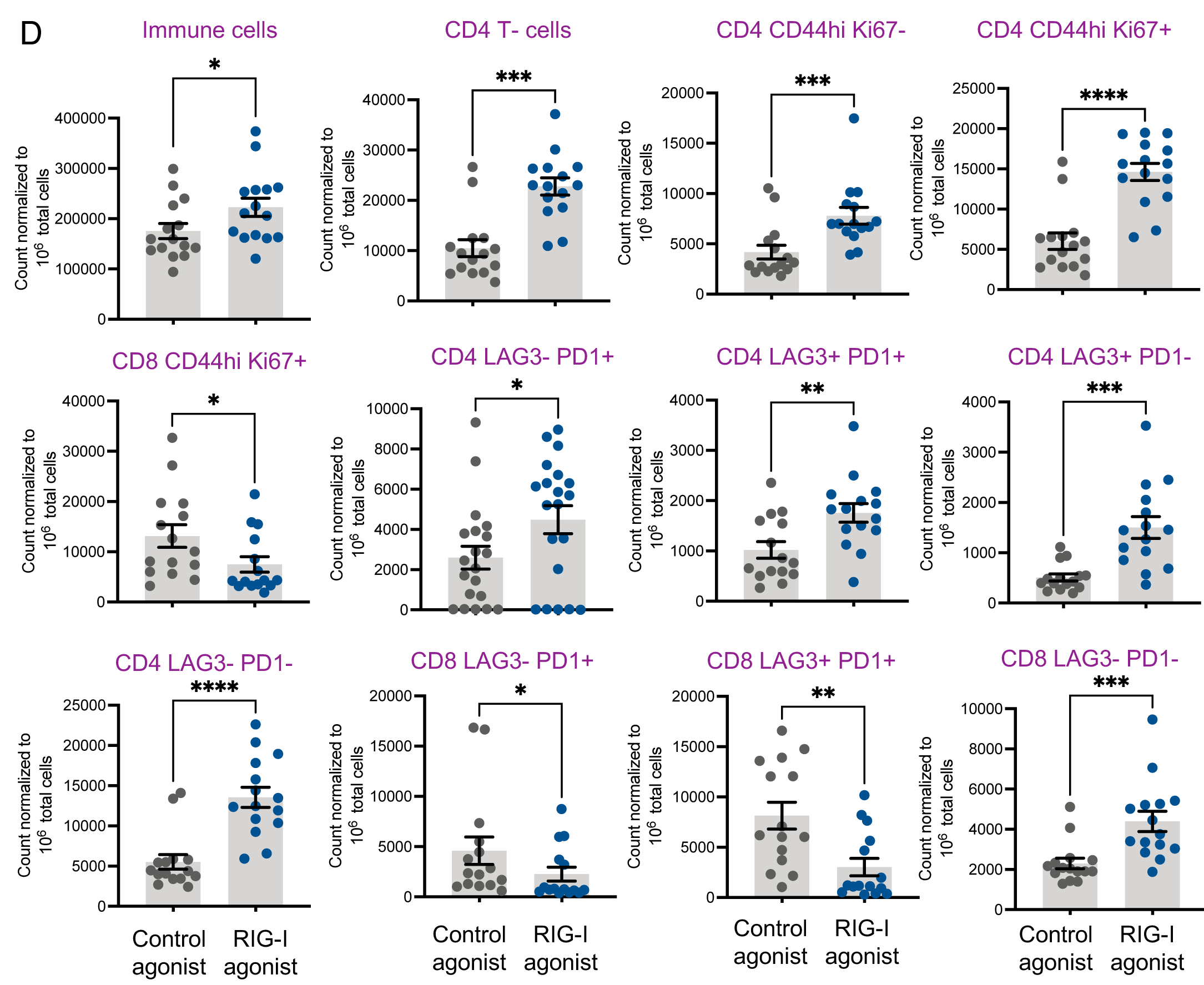
Tumor cell intrinsic RIG-I activation reprograms the molecular and cellular tumor immune microenvironment *in vivo*. (A) Nanostring Immune profiling of MC38 or CT26 tumors at endpoint (B). C-D) Representative multicolor flow cytometry data from the C) MC38 or D) CT26 tumor experiment described in Figure 3 at endpoint. Each dot represents an individual tumor. P-values from two-tailed Student’s T-test or Mann-Whitney U-test. Only the most significant cell types are depicted

### RIG-I activation correlates with immune checkpoint expression and provides therapeutic synergy

Our 37-gene RIG-I activation signature based on human and mouse datasets identified CD274 (PD-L1) as a robust RIG-I induced gene. Our RNAseq data indicated that RIG-I activation induced the transcription of specific immune checkpoints – PD-L1, Galectin-9 and LAG-3 (Figure 6A). Analysis of TCGA datasets highlighted a strong correlation between RIG-I and PD- L1, Galectin-9, LAG-3 and TIM-3, the receptor for Galectin-9 (Figure 6B). We confirmed that activation of RIG-I in tumor cells induced the cell surface expression of PD-L1 and Galectin-9 (Figure 6C-D). Using the KM plotter tool [16], we found that across all cancers, expression of RIG-I alone was a positive prognostic factor for patient survival response to immune checkpoint blockade, especially PD-L1/PD-1 antagonists (Figure 6E). Therefore, we tested whether treatment with an anti-PD-L1 antibody will enhance the effects of tumor intrinsic RIG-I. Indeed, we found that the addition of anti-PD-L1 resulted in substantial reduction of tumor growth in CT26 tumors compared to treatments with either RIG-I alone or anti-PD-L1 alone (Figure 6F-G).

**Figure 6.**
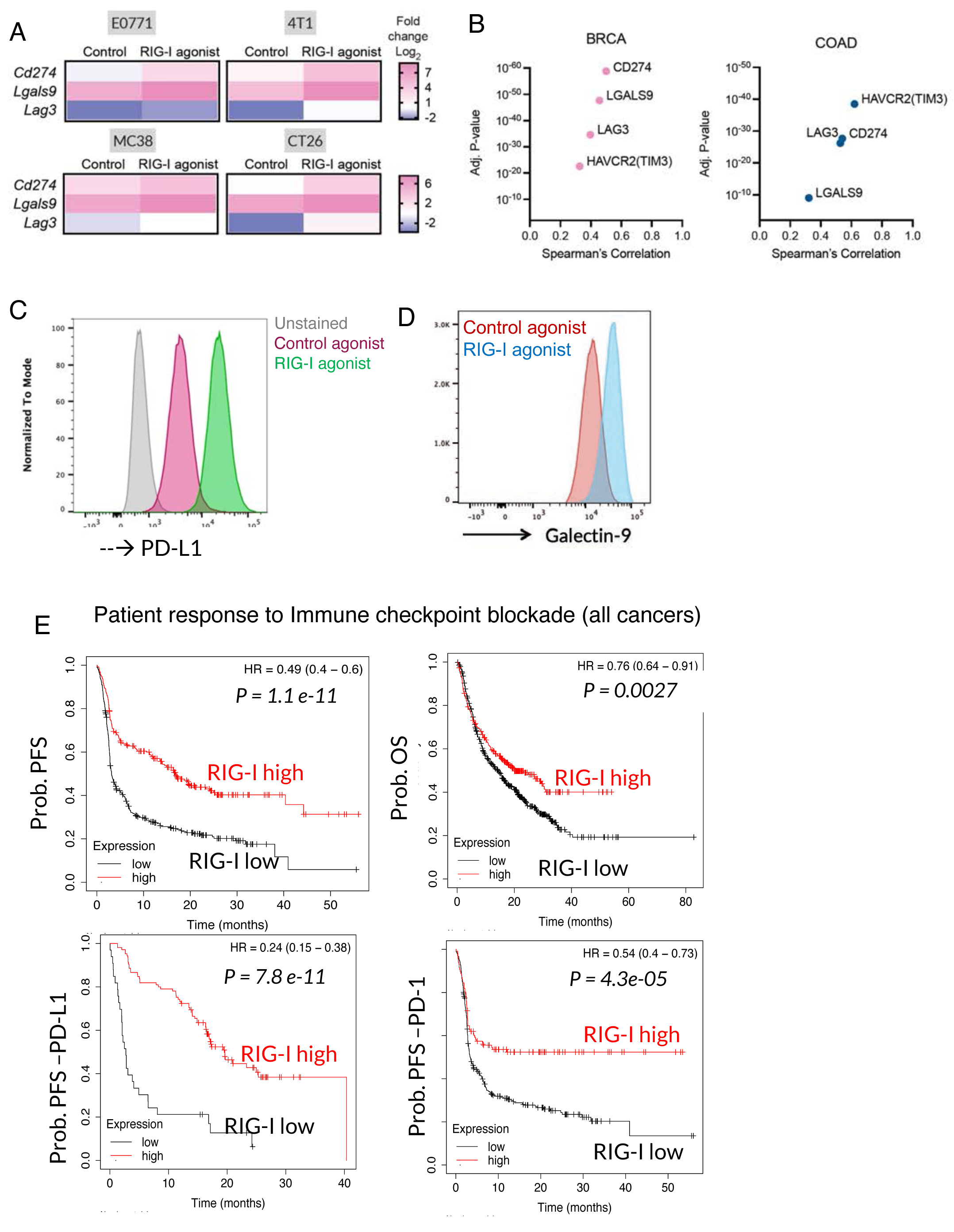
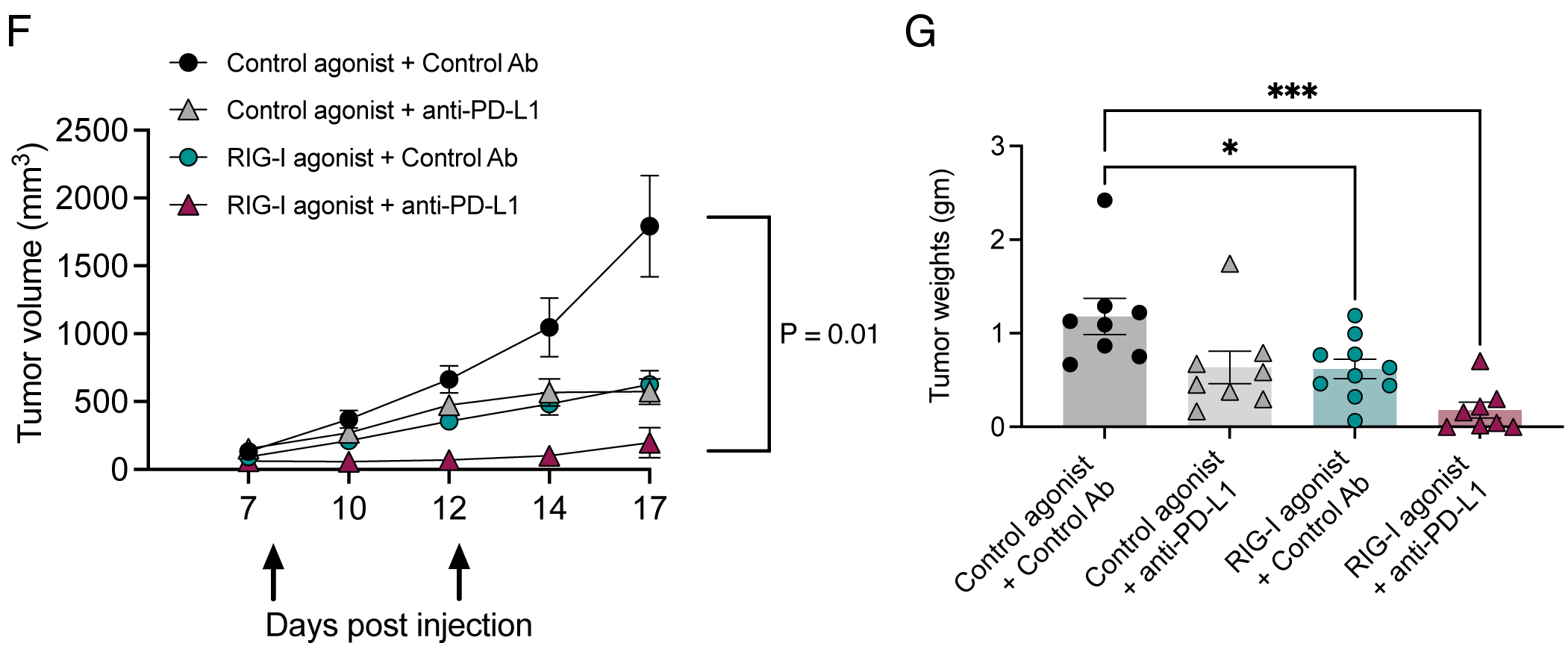
PD-L1 blockade enhances efficacy of tumor intrinsic RIG-I activation. A) Significant RIG-I (DDX58) correlated immune checkpoints in BRCA and COAD. B) RNAseq data showing induction of immune checkpoint transcripts at 24h in RIG-I agonist transfected compared to controls in the indicated cell lines. C-D) Flow cytometry validation of cell surface expression of two checkpoints PD-L1 and Galectin-9 identified in RNAseq from E0771 cells. E) Correlations between DDX58 (RIG-I) expression and responses to immune checkpoint targeted therapies in TCGA datasets across cancer types (Both CTLA4 and PD1 pathway antibodies are included for PFS and OS) analyzed via kmplot.com. The parental datasets are described in [16]. P-values from log rank test. F) Tumor kinetics and G) Tumor weights for CT26 tumor growth in Balb/C mice with indicated treatment groups. Arrows depict anti-PD-L1 injections. P-values from ANOVA with post-hoc Tukey test. Only significant comparisons are highlighted *P<0.05, **P<0.01, ***P<0.001

Taken together, our data indicates that a) RIG-I correlates with overall survival in human cancer b) immune cell landscape in RIG-I high immune tumors do not indicate robust effector activity in CRC and c) an H1N1 derived RIG-I agonist enhances tumor cell death and increases a cellular and molecular anti-tumor responses in preclinical models including immune checkpoints. Our data also highlights the utility of combining immune checkpoint, especially PD-L1 antagonists with the RIG-I agonist in preclinical colorectal cancer models. These observations suggest a complex and distinct tumor cell intrinsic role of RIG-I which may drive cell death and a potential immunological adjuvant effect where RIG-I activation enhances cellular and molecular responses in the tumor.

## Discussion

Cellular responses to non-self and occasionally, self, nucleic acids, is a potent driver of immune responses and inflammation in tissues. RIG-I recognizes cytosolic short dsRNA and plays a major role in antiviral responses [4, 17] and is known to drive a programmed cell death pathway [18]. Physiologic responses to radio-/chemo-therapy in cancers have been shown to converge on an antiviral program in the recruitment of the RIG-I-like receptors (RLR) pathway by a ncRNA-dependent activation of RIG-I [19]. This example highlights how self RNA can activate RIG-I signaling. Several studies have documented activation of RIG-I using synthetic RNA agonists results in robust anti-tumor responses across different preclinical models including pancreatic cancer, breast cancer, melanoma, leukemia and a phase I clinical trial that demonstrated safety of an RNA agonist [7-11, 20]. These observations suggest that activating RIG-I may generate durable therapeutic responses in cancer via type I interferon pathway mediated activation of immune responses.

We show here that tumor cell intrinsic activation of RIG-I using a hairpin RNA agonist derived from the H1N1 influenza virus can trigger cytotoxic effects, decrease tumor growth in vivo and robustly elicit anti-tumor immune responses. Our analysis of human tumor datasets (Fig 1B) identified a subset of cancers where RIG-I expression correlated with better overall survival. However, the comparison of molecular signatures (Fig 1C) highlighted a positive correlation across several immune signature RNAs in almost all the TCGA cancer types. We explored this further with immune profiling data from a cohort of CRC samples at OHSU and comparisons to the TCGA COAD datasets. While the molecular signatures of DDX58 high tumors revealed classic type I interferon genes such as Mx1 and ISG15, we also found members of complement and markers of T-cell exhaustion such as LAG3. We found these gene signatures were concordant between our data and the TCGA datasets. We hypothesized that activation of RIG-I using an agonist in the tumor cells will not only enhance the type I interferon signatures but also elicit cellular anti-tumor responses in CRC.

Our data from the *in vitro* experiments across tumor cell lines suggest that this new RIG-I agonist drives cell death. Gene set enrichment analyses show that in these cell lines, in addition to the interferon-α and γ response signatures, Myc targets, UPR, cell cycle checkpoints and apoptosis signatures are induced by RIG-I agonist treatment. Interestingly, there are tumor type and cell line intrinsic differences both in the magnitude and specific pathways that are activated by the RIG-I agonist. Since we had multiple mouse tumor cell line datasets and the human COAD dataset, we derived a 37-gene RIG-I activation signatures in CRC that included markers of immune activation, chemokines such as CXCL11 and the immune checkpoint PD-L1 (CD274). Given the immune and inflammatory genes, this gene signature could be a useful biomarker for prediction of responses to immunotherapy. It remains to be seen whether this RIG-I signature is unique to CRC or conserved across other tumor types.

Our preclinical tumor model studies show that ex-vivo activation of RIG-I using this agonist is sufficient to decrease tumor burden in three different tumor cell lines. Given our hypothesis derived from human CRC samples, we chose to focus our immune profiling efforts on two different well-defined preclinical CRC models -the MC38 in C57BL/6 mice and the CT26 in Balb/C mice. We found that while the RIG-I activated MC38 tumors had increase in dendritic cells, especially cDC1 and NK cells, the CT26 tumors displayed a more varied cellular immune response. We observed significant increases in antigen-experienced memory CD4 T-cells both resting and proliferative. In contrast, there were fewer proliferating memory CD8 T-cells.

However immune exhaustion as measured by PD1 and LAG3 on both CD4 and CD8 compartments was significantly decreased in the RIG-I activated tumors. Interestingly, the same CT26 tumor cells resulted in smaller tumors in immune deficient athymic nude mice with only NK cells being a significantly enriched population in these tumors. However, the effect of the RIG-I agonist disappeared in NSG mice suggesting that the immune microenvironment is still critical for tumor cell intrinsic RIG-I to effectively slow down tumor growth.

Tumor cell intrinsic activation of nucleic acid sensors can potentiate several cell death pathways. RIG-I signaling has been shown to directly promote cell killing through multiple mechanisms including intrinsic apoptosis[21, 22], extrinsic apoptosis [23], and pyroptosis [20] in diverse cell types. Our work indicates that in both colorectal and breast cancer cells, this RIG-I agonist is a strong driver of apoptosis and tumor cell death is the likely mechanism of action for the observed efficacy in the preclinical tumor models. Although we have not evaluated other modes of cell death, it is possible that more than one cell death pathway contributes to the biological consequences of RIG-I activation in these cells. In fact, our RNAseq analysis suggests in addition to cell death, RIG-I activation also induces genes responsible for cell cycle arrest.

Our immune profiling studies highlight differential effects on immune responses depending on the tumor models and background mouse strains. While we observed a few common themes such as infiltration of total lymphocytes and NK cells across tumor models, there are also model specific immune changes. For instance, the decrease in immune exhaustion and increases in the CD4 memory T-cells in the CT26 model presents opportunities for augmenting the activity of RIG-I with immunotherapy approaches.

Finally, we observed induction of immune checkpoints in our tumor cell lines in vitro and in vivo and identified strong correlations in the TCGA datasets between RIG-I and PD-L1, LAG3, Galectin-9 and its receptor TIM-3. RIG-I expression alone was a robust predictor of responses to immune checkpoint antagonists across several cancers in a publicly available cohort of immunotherapy trials. Given the considerable therapeutic interest in these pathways our observation that addition of anti-PD-L1 strongly augments the tumor intrinsic role of RIG-I opens the door to additional studies on delivering RIG-I to tumor cells in vivo, optimizing the best checkpoint to target and comparison of timing of RIG-I activation vs checkpoint blockade. Further, we wish to highlight the potential for RIG-I activated tumors to serve as autologous vaccines.

In summary, we show here that tumor cell intrinsic activation of RIG-I signaling by a novel RNA agonist might be sufficient to exert a significant anti-tumor effect and enhance the immune microenvironment. Our datasets also identify several potentially druggable opportunities in the RIG-I inflamed immune microenvironment of colorectal cancers.

## Acknowledgements

This work was supported by funding from NHLBI to S.A. (R01 HL137779 and R01 HL143803). A.B is supported by a training grant from NIGMS 5T32GM142619-02 and funds from the Knight Cancer Institute.

## Author Contributions

E. F-B, S.K. and S.A designed experiments, E. F-B, S.K., A.B., R.R., S.A performed experiments, analyzed the data, E. F-B and S.A wrote the manuscript.

## Figure Legends

**Supplementary Figure 1.**
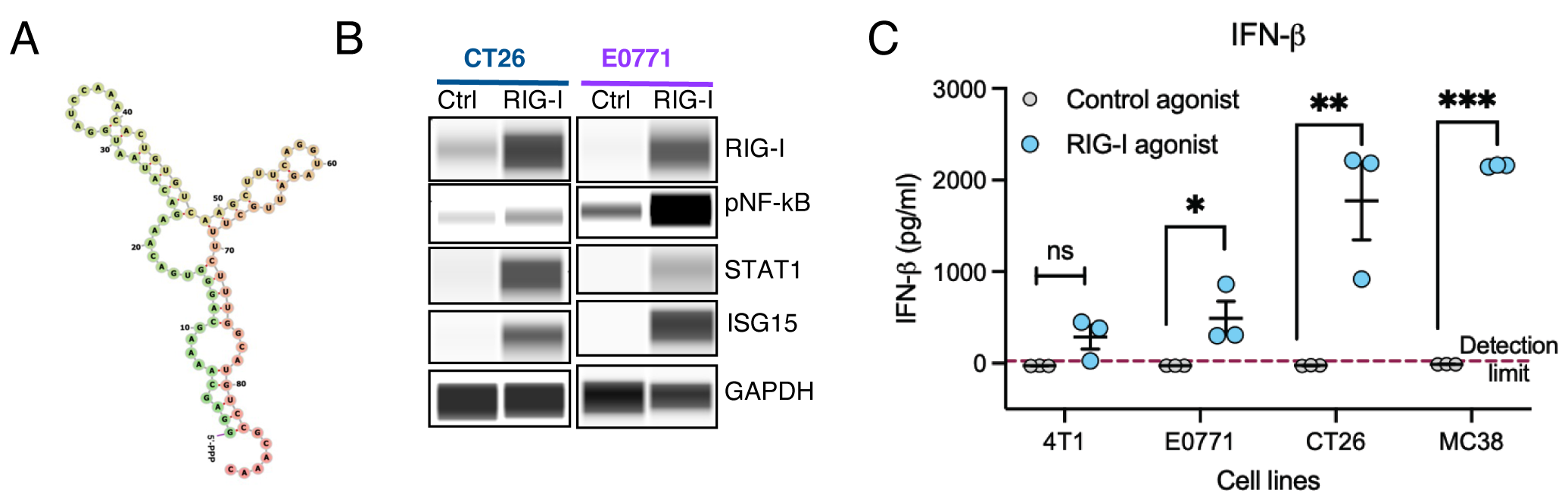
Potent and specific RIG-I activation by an agonist from H1N1 influenza virus. A) Predicted structure of the 5’-triphosphate hairpin RIG-I agonist. B) Control agonist or RIG-I agonist was transfected into CT26 or E0771 tumor cells and activation of downstream RIG-1 signaling pathways was measured by western blot of the indicated proteins using a Simple Wes assay. C) ELISA for IFN-! from cell culture supernatants 24h post transfection. * P<0.05, ** P<0.01, *** P<0.001, ****P<0.0001 using ANOVA with post-hoc Sidak’s test

**Supplementary Figure 2.**
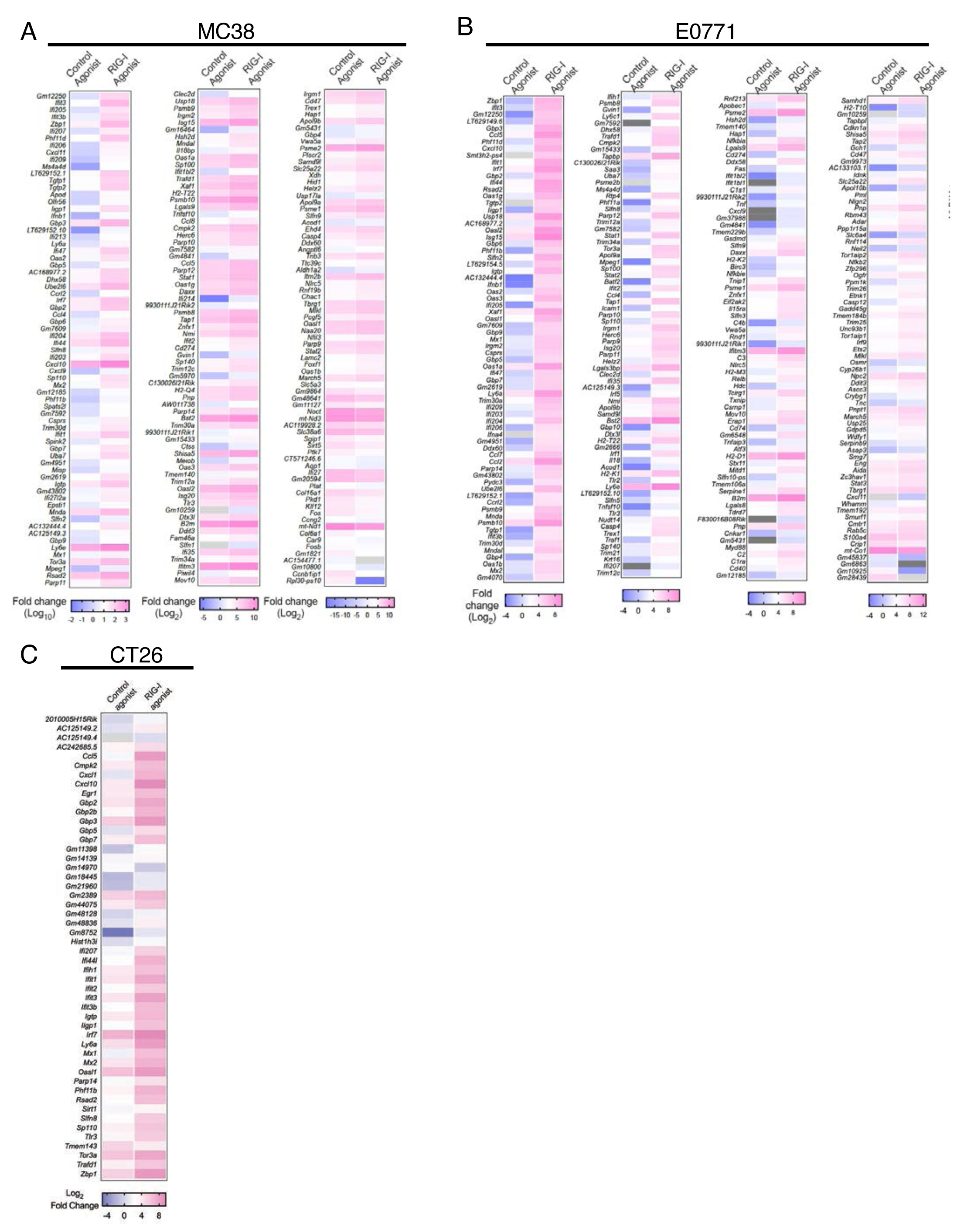
Gene expression signatures in tumor cell activation of RIG-I. Indicated cell lines were transfected with either a control agonist or a RIG-I agonist. RNA was extracted 24h later and RNA sequencing was performed. Heatmap depicts significant (Adj P-value <0.05) changes in gene expression. Median fold change of two biological replicates is shown.

**Supplementary Figure 3.**
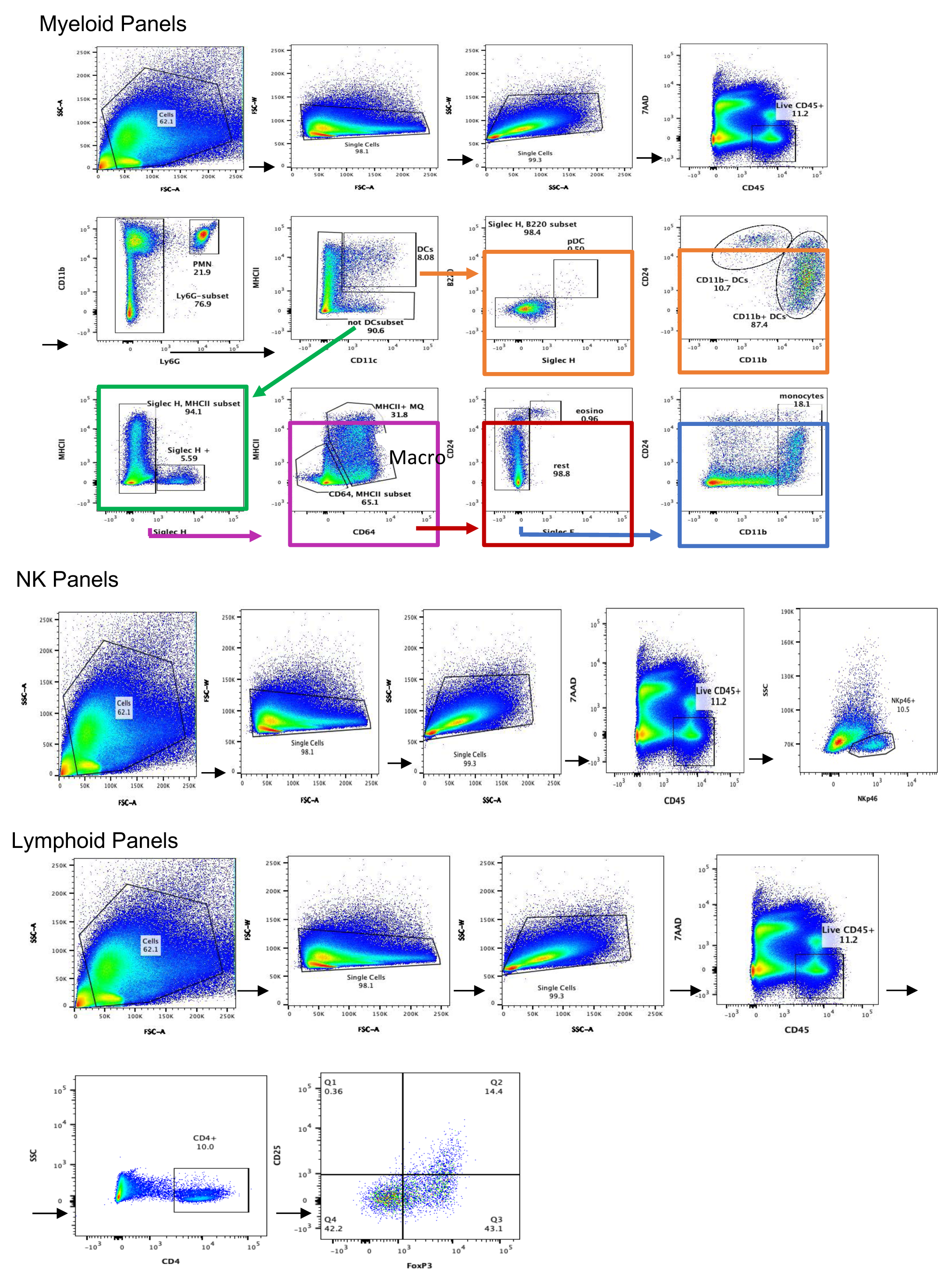
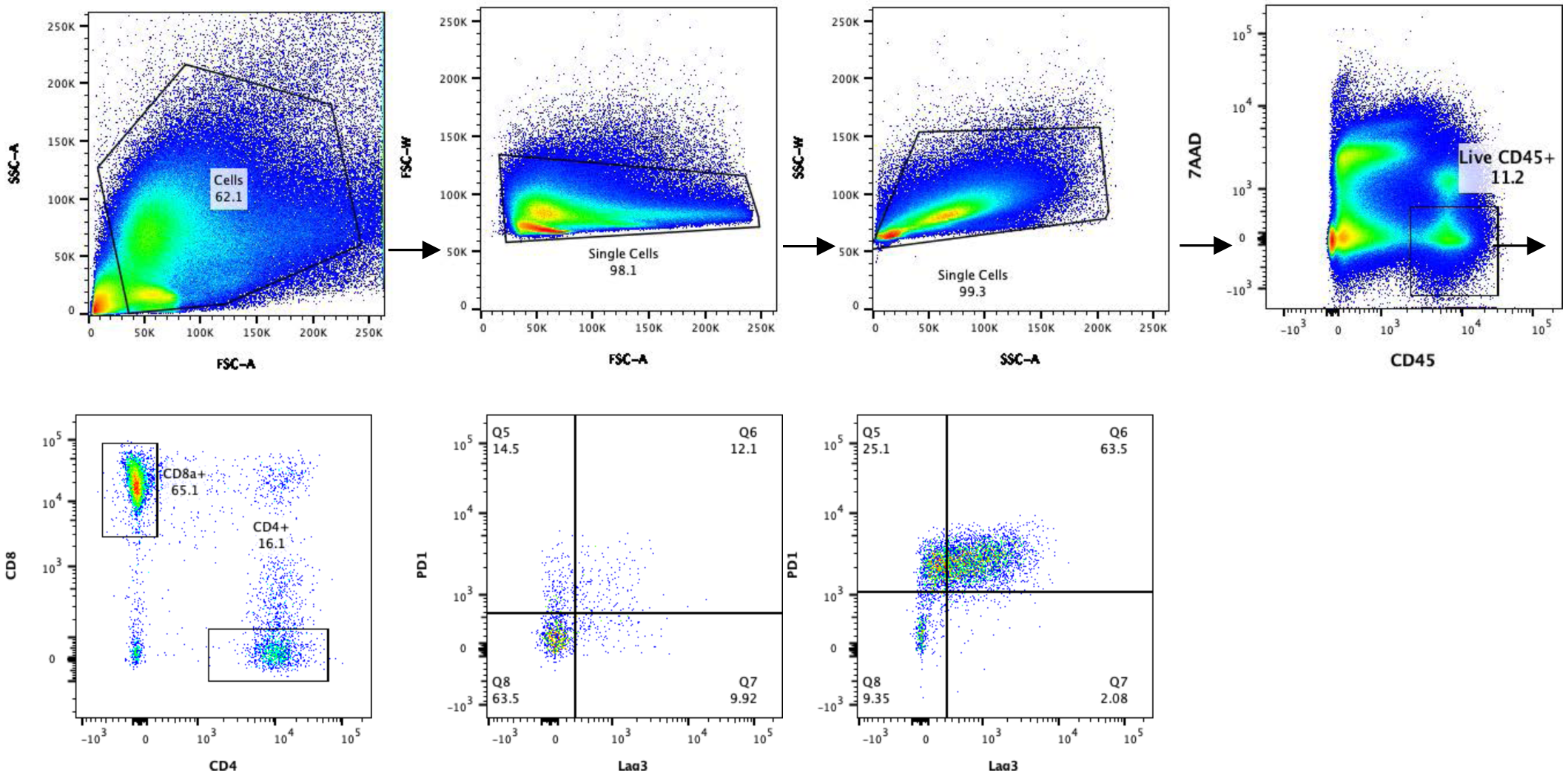
Gating strategies for multicolor flow cytometry panels.

**Supplementary Figure 4.**
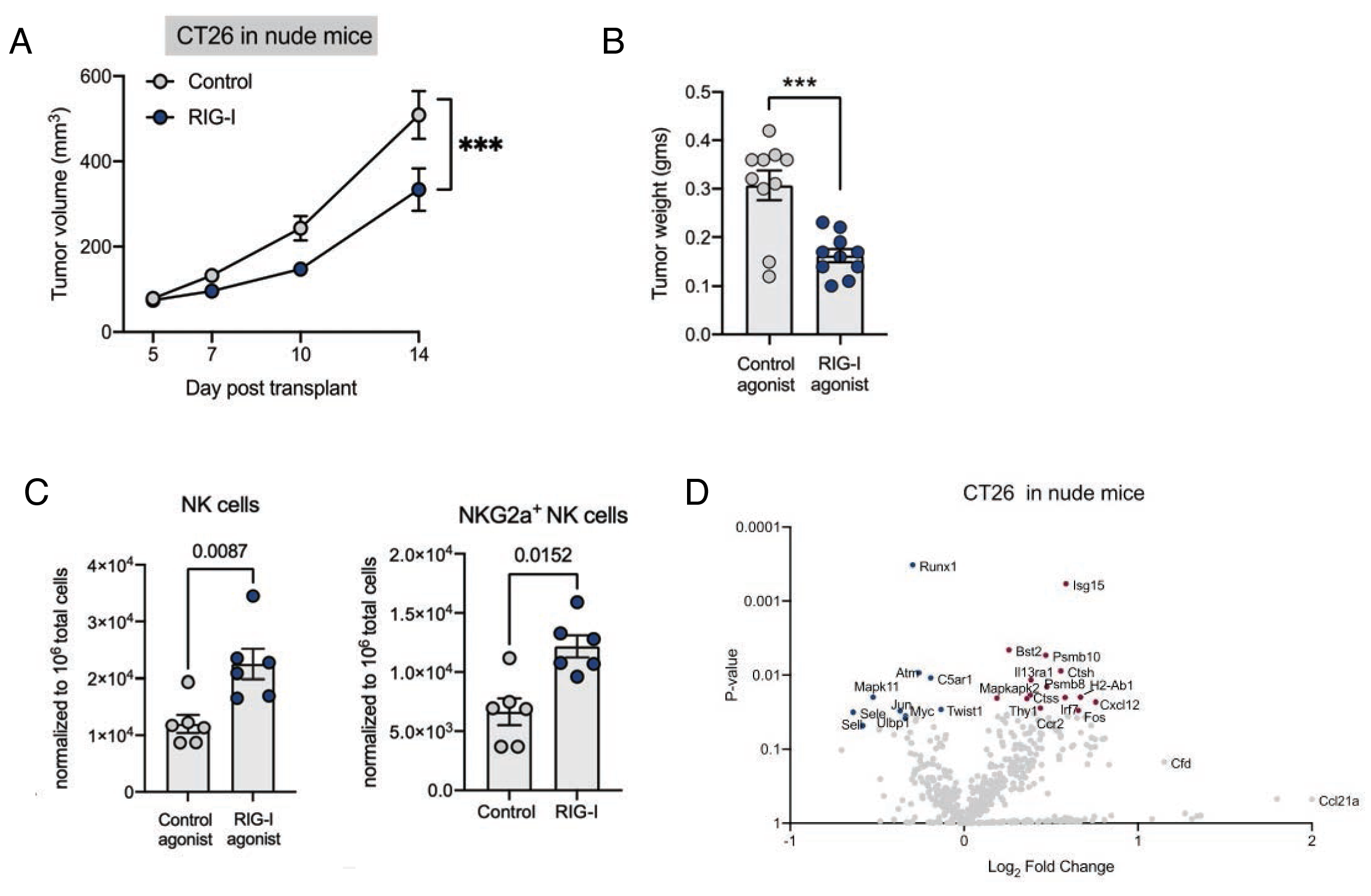
Tumor cell intrinsic RIG-I activation decreases tumor growth in athymic nude mice with modest impact on the immune microenvironment. (A) Tumor volumes (B) Tumor Weights (C) Representative multicolor flow cytometry data and (D) Nanostring Immune profiling from CT26 tumors in Balb/C nude mice. P-values from two-tailed Student’s T-test or Mann-Whitney U-test. Only the most significant cell types are depicted.

